# Quantification and image-derived phenotyping of retinal ganglion cell nuclei in the *nee* mouse model of congenital glaucoma

**DOI:** 10.1101/2021.04.13.439698

**Authors:** Carly J. van der Heide, Kacie J. Meyer, Adam Hedberg-Buenz, Danielle Pellack, Nicholas Pomernackas, Hannah E. Mercer, Michael G. Anderson

## Abstract

The *nee* mouse model exhibits characteristic features of congenital glaucoma, a common cause of childhood blindness. The current study of *nee* mice had two components. First, the time course of neurodegeneration in *nee* retinal flat-mounts was studied over time using a retinal ganglion cell (RGC)-marker, BRN3A; a pan-nuclear marker, TO-PRO-3; and H&E staining. Based on segmentation of nuclei using ImageJ and RetFM-J, this analysis identified a rapid loss of BRN3A^+^ nuclei from 4–15 weeks of age, with the first statistically significant difference in average density compared to age-matched controls detected in 8-week-old cohorts (49% reduction in *nee*). Consistent with a model of glaucoma, no reductions in BRN3A^−^ nuclei were detected, but the combined analysis indicated that some RGCs lost BRN3A marker expression prior to actual cell loss. These results have a practical application in the design of experiments using *nee* mice to study mechanisms or potential therapies for congenital glaucoma. The second component of the study pertains to a discovery-based analysis of the large amount of image data with 748,782 segmented retinal nuclei. Using the automatedly collected region of interest feature data captured by ImageJ, we tested whether RGC density of glaucomatous mice was significantly correlated to average nuclear area, perimeter, Feret diameter, or MinFeret diameter. These results pointed to two events influencing nuclear size. For variations in RGC density above approximately 3,000 nuclei/mm^2^ apparent spreading was observed, in which BRN3A^−^ nuclei—regardless of genotype—became slightly larger as RGC density decreased. This same spreading occurred in BRN3A^+^ nuclei of wild-type mice. For variation in RGC density below 3,000 nuclei/mm^2^, which only occurred in glaucomatous *nee* mutants, BRN3A^+^ nuclei became smaller as disease was progressively severe. These observations have relevance to defining RGCs of relatively higher sensitivity to glaucomatous cell death and the nuclear dynamics occurring during their demise.

## 1. Introduction

Mice homozygous for the spontaneously arising *Sh3pxd2b*^*nee*^ mutation (i.e. *nee* mice) exhibit anterior segment dysgenesis and features of severe congenital glaucoma. In two different genetic backgrounds, the original C57BL/10-related and a congenic C57BL/6J background, *nee* mice develop high intraocular pressure (IOP) followed by retinal ganglion cell (RGC) loss which is profound by 3 months of age (Daniel et al., 2019; Mao et al., 2011). Along with congenital glaucoma, *nee* mice exhibit craniofacial, cardiac and skeletal abnormalities that recapitulate Frank-Ter Haar syndrome (Mao et al., 2009), a rare human disease caused by recessive mutations in the homologous gene, *SH3PXD2B* (Iqbal et al., 2010). Thus, *nee* mice directly model a known cause of glaucoma in humans. The manifestation of glaucoma in *nee* mice is also notable among animal models of glaucoma because the disease is spontaneous, severe, and fully penetrant (Mao et al., 2011), which are all experimental advantages.

While RGC loss in *nee* mice is known to be relatively severe and early onset, refining the precise time-course of RGC loss in *nee* could uncover new insights and is a prerequisite needed to inform the design of many future experiments, such as utilizing the *nee* model in tests of neuroprotective agents. To achieve this, we have characterized the progression of RGC death in young *nee* mice using ImageJ in a high-throughput image analysis technique that quantified thousands of cell nuclei in the inner retina of each mouse retina. In addition to providing counts of RGC nuclei, our image analysis data included information about the size and shape of nuclei. In a discovery-based approach, we used these data to explore whether RGC density of glaucomatous mice was significantly correlated to average nuclear area, perimeter, Feret diameter, or MinFeret diameter. Combined, the results identify ages past 8 weeks as having statistically detectable differences in RGC density, identify a caveat regarding loss of RGC marker expression prior to RGC cell loss in some cells of *nee* mice, and point to the possible ways in which automatedly collected image data can be used to discover new disease-associated phenotypes.

## 2. Materials and methods

### 2.1 Animal Husbandry

Generation of N10 congenic B6.*Sh3pxd2b*^*nee*^ mice has been previously described (Daniel et al., 2019). The *nee* strain was maintained in heterozygote X heterozygote intercrosses, with littermates homozygous for the mutant (abbreviated throughout as “*nee*”) or wild-type (abbreviated throughout as “WT”) alleles primarily utilized in this study. As indicated in the text, the SD-OCT imaging study utilized both WT and heterozygous mice (referred to as “non-mutant”, abbreviated throughout as “non-mut”) to increase the sample size of controls. Mice were housed at the University of Iowa Animal Research Animal Facility, where they were maintained on a 4% fat NIH 31 diet provided ad libitum, housed in cages containing dry bedding (Cellu-dri; Shepherd Specialty Papers, Kalamazoo, MI), and kept in a 21ºC environment with a 12-h light: 12-h dark cycle. All experiments included both male and female mice. All mice were treated in accordance with the Association for Research in Vision and Ophthalmology Statement for the Use of Animals in Ophthalmic and Vision Research. All experimental protocols were approved by the Institutional Animal Care and Use Committee of the University of Iowa.

### 2.2 Preparation of retinal flat-mounts for RGC quantification

Mice were euthanized using CO_2_ asphyxiation followed by cervical dislocation and eyes were immediately placed in cold 4% paraformaldehyde for 10 minutes. Posterior cups were dissected in PBS and replaced in cold 4% paraformaldehyde for 1 hour. Posterior cups were washed with PBS, and permeabilized with 0.3% Triton-X 100 in PBS (PBST) overnight at 37°C. All subsequent steps were carried out at room temperature unless otherwise specified. Posterior cups were incubated in 3% H_2_O_2_-1% NaHPO_4_ for 2 minutes and then retinas were dissected in PBS. Retinas were replaced in 3% H_2_O_2_-1% NaHPO_4_ for 1 hour. Retinas were washed with PBST three times for 15 minutes, and then further permeabilized with PBST at -80°C for 15 minutes. Retinas were washed again with PBST twice for 15 minutes, then blocked with 2% normal donkey serum in PBST overnight at 4°C. Retinas were then incubated in primary antibody (additional details noted in paragraph below) in PBS with 2% normal donkey serum, 1% Triton-X 100, and 1% DMSO (primary buffer) for two overnights at 4°C. Retinas were washed for 5 minutes in PBS with 2% Triton X-100, then washed further with PBST four times for 15 minutes. Then retinas were incubated in secondary antibody (additional details below) diluted in PBS with 5% normal donkey serum, 1% Triton-X 100, and 1% DMSO (secondary buffer). Retinas were washed twice for 15 minutes in PBST, then incubated in 1 μM TO-PRO-3 (T3605; Thermo Fisher Scientific, Waltham, MA) in PBS for 15 minutes, and washed again with PBST three times for 15 minutes. Retinas were then washed briefly with PBS and mounted using Aqua-Mount (Lerner, Pittsburgh, PA). Individual retinas that were studied by both immunolabeling and hematoxylin/eosin (H&E) staining were labeled as described above, followed by soaking off coverslips by submersion of the slide in distilled water. H&E staining was performed under the following conditions: hematoxylin (1 minute), rinse in double-distilled water, acid alcohol (3 seconds), rinse in water, 80% ethanol (1 minute), eosin (2 seconds), 95% ethanol (30 seconds), 100% ethanol (30 seconds), and xylenes (twice for 30 seconds each time). MM 24 mounting media (Leica, Wetzlar, Germany) was used to apply coverslips to slides.

Different antibody combinations were applied to the protocol (as described above) for two distinct purposes. For the purpose of RGC quantification, retinas were incubated in a polyclonal goat anti-BRN3A antibody (C-20, 1:200; Santa Cruz Biotechnology, Dallas, TX) diluted in primary buffer for two overnights at 4°C followed by incubation in a donkey anti-goat Alexa 488-conjugated secondary antibody (A11055, 1:200; Life Technologies, Madison, WI) diluted in secondary buffer for three hours at room temperature. For the purpose of testing for potential marker-specific effects in quantifying RGCs, retinas were incubated in the anti-BRN3A and a polyclonal rabbit anti-RBPMS antibody (GTX118619, 1:1,000; GeneTex, Irvine, CA) diluted in primary buffer for two and four overnights at 4°C, respectively. Retinas underwent two sequential treatments in secondary antibodies diluted in secondary buffer, including an incubation in the donkey anti-goat Alexa 488-conjugated secondary antibody as indicated above and then in a donkey anti-rabbit Alexa 546-conjugated secondary antibody (A10040, 1:500; Life Technologies, Madison, WI) overnight at 4°C.

### 2.3 Microscopy and quantification of nuclei in the inner retina

Immunolabeled retinas were photographed using a Zeiss LSM710 confocal microscope (Carl Zeiss, Oberkochen, Germany). Images were collected at ∼538X total magnification (1024×1024 px, 0.18 mm^2^ image area) from non-overlapping fields at each of three zones of eccentricity (n = 12 images total: 4 central, 4 mid-peripheral, 4 peripheral). Photos were captured in a single plane of focus at the level of the RGC layer. Images were processed in ImageJ with the Enhance Local Contrast (CLAHE) process (blocksize 125, histogram bins 256, maximum set to 3 for BRN3A-labeled images and maximum set to 10 for TO-PRO3-labeled images), the Subtract Background tool with rolling ball radius set to 35 pixels for BRN3A and 20 pixels for TO-PRO-3, followed by the Smooth tool. The TO-PRO-3 images were additionally processed with Gaussian blur (sigma set to 0.5) and the Enhance contrast process with 10% saturated pixels and application of histogram equalization. Images were then converted to binary using Huang thresholding. Binary images were further processed using the Open, Watershed, and Fill Holes functions. To count BRN3A^+^ nuclei, the Analyze Particles function was applied to the BRN3A images with particle size set to 38–150 μm^2^ and circularity 0.55–1. The segmentation of BRN3A^+^ nuclei was saved as a region of interest and subtracted from the TO-PRO-3 images. Then, the Analyze Particles function was applied to TO-PRO-3 images with size set to 20–150 μm^2^ and circularity of 0.4–1 to quantify the BRN3A^−^, TO-PRO-3^+^ population. The metrics quantified with the Analyze Particles function are defined by ImageJ as follows: “Area” refers to the area of the selection in square pixels, calibrated to square microns; “Perimeter” refers to the length of the outside boundary of the selection; “Feret diameter” refers to the longest distance between any two points along the selection boundary, a.k.a. maximum caliper; “MinFeret diameter” refers to the minimum caliper. Individual retinas that were studied by both immunolabeling and H&E staining were photographed and quantified as described above, followed by H&E staining and imaging with a light microscope (BX52; Olympus, Tokyo, Japan) equipped with a digital camera (DP72; Olympus, Tokyo, Japan). H&E-stained nuclei were quantified with RetFM-J as previously described (Hedberg-Buenz et al., 2016c).

In testing for the possibility of marker-specific results, an independent set of retinas immunolabeled for both BRN3A and RBPMS were imaged as described above. Four images, including two from the peripheral and mid-peripheral zones of retina were randomly selected for manual quantification of cells immunolabeled by each marker using the “cell counter” tool of ImageJ {Schneider, 2012 #1} by an investigator blinded to the identity of each image (i.e. genotype of mouse the retina was collected from, retinal zone, etc.). All images were viewed by the investigator in single-channel format, and therefore single-marker, to ensure that cell detection and counting is done using the signal emitted by only one marker at a time. Results were expressed in terms of cell number and compared across markers using a Bland Altman plot and coefficient of determination (R^2^).

### 2.4 Optical coherence tomography

8-week-old mice received an intraperitoneal injection of a mixture of ketamine/xylazine (87.5 mg/kg Ketamine (VetaKet®, AKORN, Lake Forest, IL)/12.5 mg/kg Xylazine (Anased, Lloyd Laboratories®, Shenandoah, IA). Upon anesthesia, eyes were hydrated with balanced salt solution (BSS; Alcon Laboratories, Fort Worth, TX). To obtain retinal images, the tear film was wicked away and eyes were imaged with a Bioptigen spectral domain optical coherence tomographer (SD-OCT; Bioptigen, USA) using the mouse retinal bore. The volume intensity projection was centered on the optic nerve. Following imaging, eyes were hydrated with lubricant ophthalmic ointment (Artificial Tears, AKORN, Lake Forest, IL) and mice were provided supplemental indirect warmth for anesthesia recovery. Retinal ganglion cell complex (RGCC; inner limiting membrane to the innermost border of the inner nuclear layer) thickness was measured from retinal images within the Bioptigen InVivoVue Clinic software four times per eye using vertical angle-locked B-scan calipers.

### 2.5 Optic nerve collection, histology, and quantification

8-week-old mice were euthanized by carbon dioxide inhalation with death confirmed by cervical spine dislocation. Optic nerves were collected, processed for histology, stained, imaged, and quantified as previously described (Mao et al., 2011; Trantow et al., 2009). Mouse heads were fixed in half-strength Karnovsky’s fixative for a minimum of 24 hours. Nerves were then dissected and embedded in resin. Histologic sections were cut using an ultramicrotome (UC6, Leica, Wetzler, German) equipped with a diamond knife (Histo, Diatome, Hatfield, PA, USA), stained with paraphenylenediamine (PPD; (Anderson et al., 2005)), and imaged using a light microscope (BX52; Olympus, Tokyo, Japan) equipped with a camera (DP72; Olympus, Tokyo, Japan) and corresponding software (CellSens; Olympus, Tokyo, Japan). Cross-sectional area of the optic nerve was measured using ImageJ. Axons were counted from 6–10 non-overlapping fields at 1000X magnification representing 10% of the optic nerve cross-sectional area. The total number of axons for each nerve was estimated by multiplying the sum of the axons in the counted fields by 10.

### 2.6 Statistics

Nuclear quantification data were analyzed using 2-way ANOVA and corrected for multiple comparisons using Sidak’s multiple comparisons test with a single pooled variance. Correlations were analyzed for significance using linear regression. Data comparing nuclear area and retinal area were analyzed by multiple two-tailed Student’s *t*-tests and corrected for multiple comparisons using the two-stage step-up method of Benjamini, Krieger and Yekutieli. The power calculations noted in the Discussion was performed using the Piface Java applet by Russell V. Lenth (Lenth, 2006) with an analysis for two-sample *t*-test (pooled or Satterthwaite). For the BRN3A^+^ cell density, the power calculation assumptions entered are sigma1 = 794.2 (standard deviation of *nee* BRN3A^+^ cell density at 8 weeks old), sigma2 = 162.0 (standard deviation of WT BRN3A^+^ cell density at 8 weeks old), alpha = 0.05, true difference of means = 870.2 (50% of the BRN3A^+^ cell loss in *nee* mice at 8 weeks old compared to WT littermates), power = 0.8981, and solving for sample size. For the retinal ganglion cell complex (RGCC) thicknesses measured by OCT, the power calculation assumptions entered are sigma1 = 14.3 (standard deviation of *nee* RGCC thickness at 8 weeks old), sigma2 = 1.7 (standard deviation of non-mut RGCC thickness at 8 weeks old), true difference of means = 11.7 (50% of the RGCC thickness decrease in *nee* mice at 8 weeks old compared to non-mut littermates), power = 0.9019, and solving for sample size. For the optic nerve cross-sectional area, the power calculation assumptions entered are sigma1 = 0.007 (standard deviation of *nee* cross-sectional area at 8 weeks old), sigma2 = 0.026 (standard deviation of the WT cross-sectional area at 8 weeks old), alpha = 0.05, true difference of means = 0.022 (50% of the optic nerve cross-sectional area loss in *nee* mice at 8 weeks old compared to WT littermates), power = 0.9084, and solving for sample size. For the optic nerve total axon number, the power calculation assumptions entered are sigma1 = 5,325 (standard deviation of *nee* optic nerve axon number at 8 weeks old, sigma2 = 4,279 (standard deviation of WT optic nerve axon number at 8 weeks old), true difference of means = 8,884 (25% of the optic nerve axon number loss in *nee* mice at 8 weeks old compared to WT littermates), power = 0.9256, and solving for sample size. A two-tailed Student’s *t*-test was used to analyze data comparing *nee*-mutant mice vs. non-mutant mice for OCT RGCC thickness, optic nerve cross-sectional area, and total axon number.

## 3. Results

### 3.1 Severe RGC loss in nee mice by 8 weeks of age

To refine the early progression of congenital glaucoma in *nee* mice, we quantified RGC loss in *nee* and WT littermates from 4–15 weeks of age. Immunolabeling of retinal flat-mounts with an anti-BRN3A antibody showed a progressive loss of BRN3A^+^ RGCs (Fig. 1A). At 4 weeks of age, quantified data showed a significant loss of BRN3A^+^ nuclei in all retinas by 8 weeks of age (Fig. 1B), whereby *nee* mice exhibited a 49% reduction in the average density of BRN3A^+^ nuclei (1,801.2 ± 794.2 nuclei/mm^2^) compared to WT littermates (3,541.3 ± 162.0 nuclei/mm^2^; p < 0.05, 2-way ANOVA with Sidak’s multiple comparisons correction). BRN3A^+^ RGC densities in *nee* mice younger than 8 weeks of age were highly variable, with some eyes severely affected while others were not different from WT littermates. As a test for the possibility of marker-specific results, an independent group of flat-mounts were simultaneously analyzed for BRN3A and RBPMS immunolabeling (S. Fig. 1). The results again indicated a severe loss of RGCs detectable by both markers in nee mice. Comparisons of the two markers were broadly congruent with one another, though RBPMS labeled more cells that BRN3A, particularly in severe disease.

**Fig. 1.**
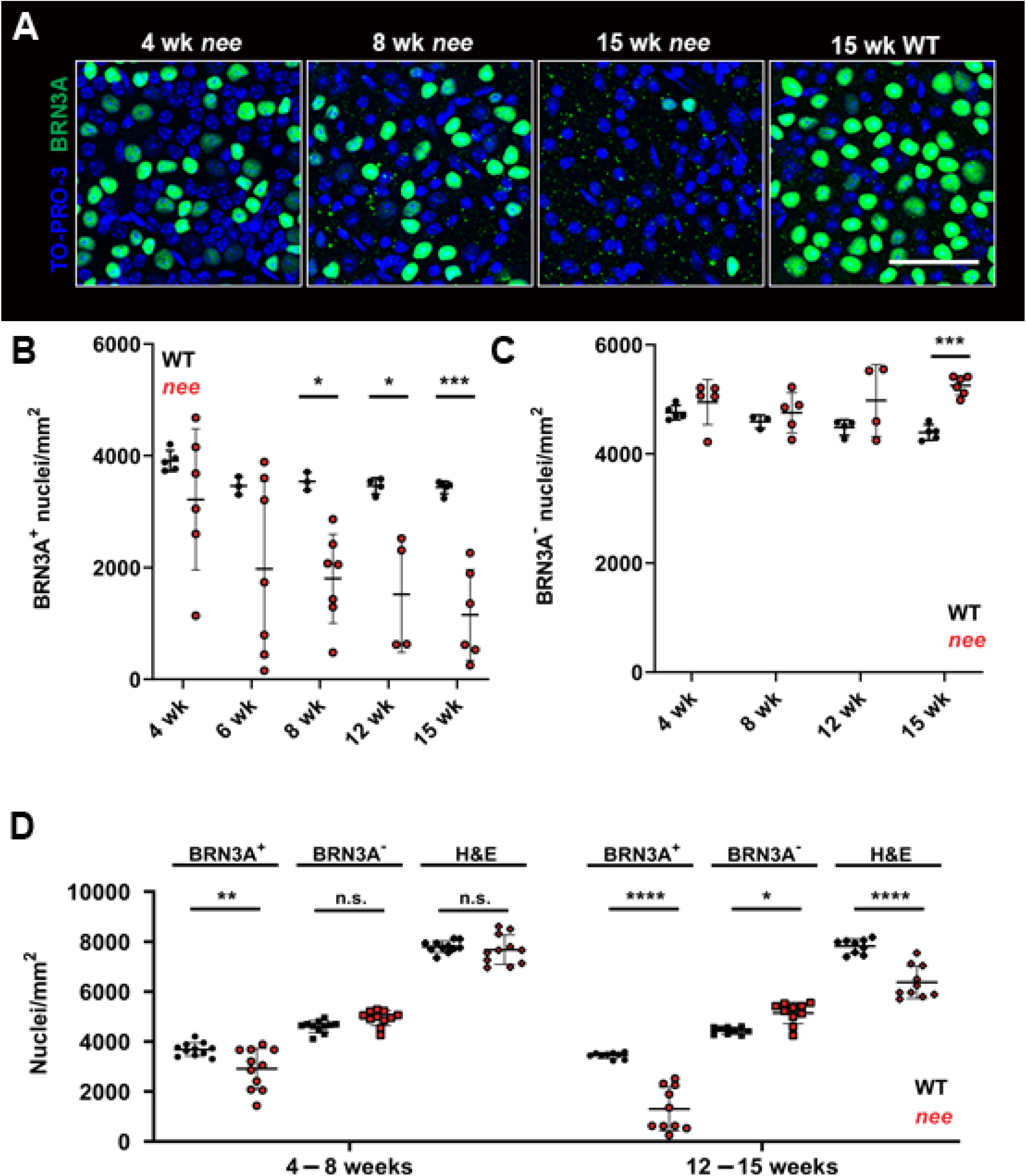
Early loss of BRN3A^+^ RGCs in *nee* mice. **(A)** Representative images showing retinal flat-mounts with BRN3A immunolabeling of RGC nuclei (*green*) and all TO-PRO-3 labeled nuclei (*blue*) in *nee* and WT mice. Confocal images taken from the peripheral region of retinas; scale bar = 50 μm. **(B)** Density plots of BRN3A^+^ RGC nuclei in *nee* (*red*) and WT (*black*) retinas showed a significant loss in RGC density in 8-week-old and older *nee* mice. Note, some eyes in the 4- and 6-week groups were severely affected. **(C)** Average density of BRN3A^−^ nuclei in *nee* retinas had an upward trend with increasing age, with a statistically significant difference at 15 weeks of age. **(D)** Plots of nuclear densities with data from B and C binned by age and additionally showing density of H&E-detected nuclei. There was no difference in density of H&E labeled nuclei between *nee* and WT retinas in the 4–8-week bin, even though these same retinas had a significant reduction in BRN3A^+^ nuclei; among *nee* retinas in the 12–15-week bin, the loss of BRN3A^+^ nuclei, off-set by an increase in BRN3A^−^ nuclei, approximated the net change in nuclei detected with H&E. n.s., not significant; *p < 0.05; **p < 0.01; ***p < 0.001; ****p < 0.0001.

To further evaluate *nee* mice as a model of glaucoma, we tested whether non-RGC cell types die in *nee* retinas by quantifying nuclei labeled with TO-PRO-3 but negative for BRN3A (BRN3A^−^). There was no reduction in the density of BRN3A^−^ nuclei of *nee* mice in any age group (Fig. 1C). Oddly, *nee* mice exhibited a 16% *higher* density of BRN3A^−^ nuclei at 15 weeks (5,253.6 ± 176.0 nuclei/mm^2^) compared to WT littermates (4,393.4 ± 142.3 nuclei/mm^2^ p = 0.0004, 2-way ANOVA with Sidak’s multiple comparisons correction). Suspecting that the increased density of BRN3A^−^ nuclei observed in *nee* mice was due to a loss of marker expression, not cell loss, these retinas were stained with H&E and nuclei of all cell types were counted (Hedberg-Buenz et al., 2016c). In *nee* mice aged 4–8 weeks, there was a 21% reduction in the density of BRN3A^+^ nuclei (2,909 ± 817 nuclei/mm^2^) compared to WT (3,686 ± 265 nuclei/mm^2^; p < 0.01; 2-way ANOVA with Sidak’s multiple comparisons correction), but the density of total nuclei stained by H&E exhibited no corresponding cell loss (7,675 ± 587 in *nee* vs 7,797 ± 228 nuclei/mm^2^ in WT; Fig. 1D), a discrepancy that suggests loss of only BRN3A immunoreactivity. Slightly older *nee* mice (12–15 weeks old) exhibited a 62.3% loss of BRN3A^+^ RGCs (1,299 ± 873 in *nee* vs 3,444 ± 120 nuclei/mm^2^ in WT; p < 0.0001; 2-way ANOVA with Sidak’s multiple comparisons correction) along with a 18.5% decrease in total cells stained with H&E (6,373 ± 646 in *nee* vs 7,816 ± 287 nuclei/mm^2^ in WT). Because RGCs constitute approximately 41–50% of total RGC-layer cells in C57BL/6J mice (Jeon et al., 1998; Schlamp et al., 2013), if only RGCs were lost, an 18.5% decrease in total cells would correspond to a 37– 45% decrease in RGCs. The RGC loss we detected using BRN3A immunolabeling is 17–25% higher than the H&E estimate, a difference similar to the 21% discrepancy we detected in the younger cohort. Conclusively, we estimate that ∼20% of RGCs in glaucomatous *nee* mice stop expressing BRN3A prior to actual RGC loss.

Non-invasive in vivo imaging of the retina was performed in 8-week-old glaucomatous *nee* mice using SD-OCT. Corresponding to the severe loss of RGCs, the RGC complex is significantly thinner in 8-week-old *nee* mice (n = 6) compared to non-mutant (HET and WT genotypes combined; n = 5) littermates (32.9 ± 14.3 μm vs. 56.3 ± 1.7 μm; p < 0.01; Student’s two-tailed *t*-test; Fig. 2A-C). We also observed differences in the appearance of optic nerve head depth and the peripapillary retina. Retinas of *nee* mice typically had an overall normal lamination, though 4 out of 21 retinas examined at 8 weeks of age had lamination abnormalities and/or retinal rosettes. As a complementary approach, optic nerve cross-sectional area and axon counts were quantified in 8-week-old *nee* mice (Fig. 2D-G). There was a significant reduction in both optic nerve cross-sectional area (0.050 ± 0.007 mm^2^ vs. 0.093 ± 0.26 mm^2^; p < 0.001; Student’s two-tailed *t*-test; Fig. 2F) and axon number (12,660 ± 5,326 axons vs. 48,196 ± 4,280 axons; p < 0.001; Student’s two-tailed *t*-test; Fig. 2G) in *nee* mice (n = 8) compared to WT littermates (n = 5).

**Fig. 2.**
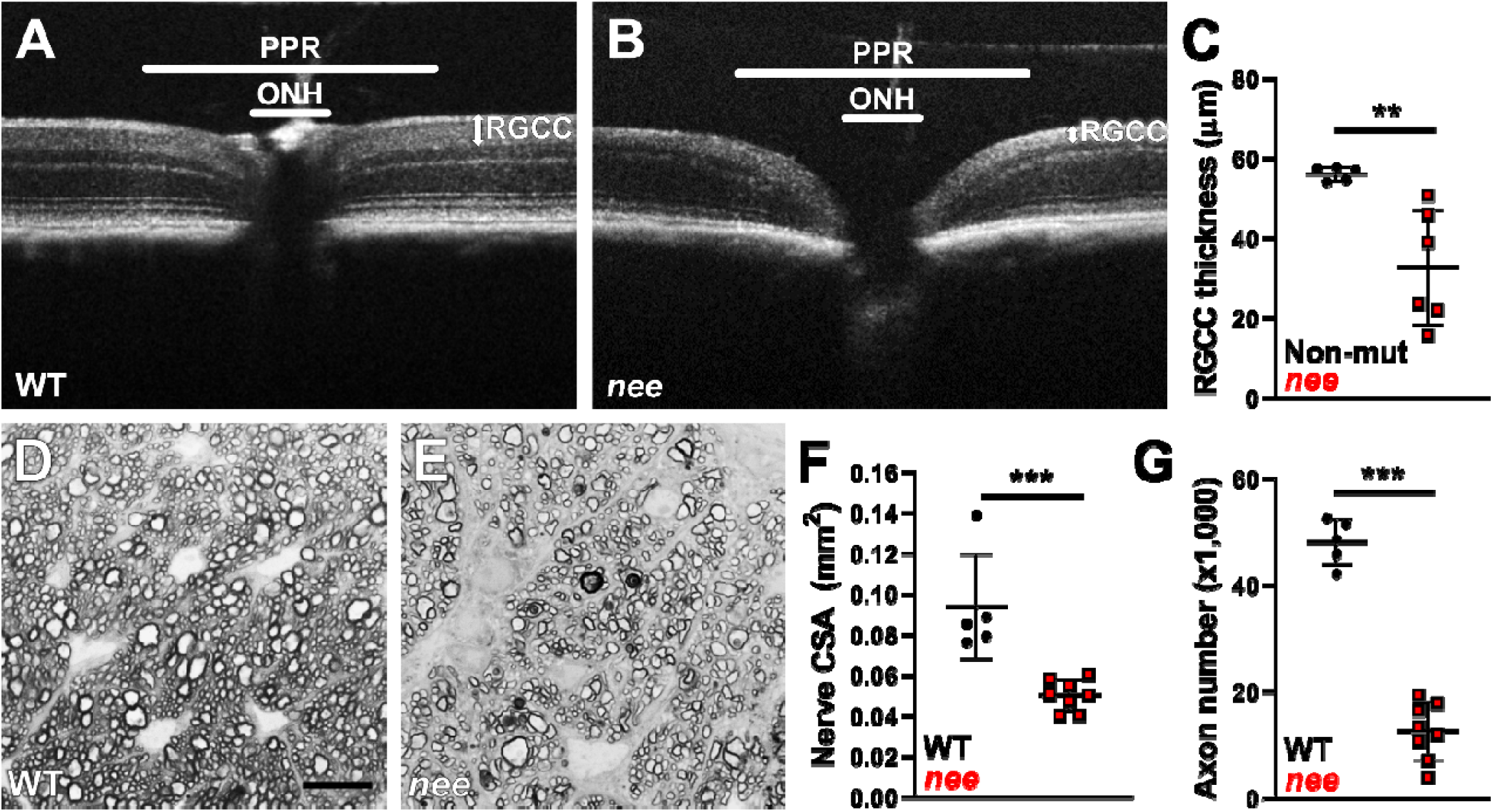
Thinning of retinal ganglion cell complex and loss of optic nerve axons in 8-week-old *nee* mice. Representative images showing in vivo retinal optical coherence tomography from 8-week-old **(A)** WT and **(B)** *nee* mice. Note differences in the optic nerve head (ONH) region, the peripapillary retina (PPR) and the retinal ganglion cell complex (RGCC). **(C)** Plot of retinal ganglion cell complex thickness showing a statistically significant difference in 8-week-old non-mutant (WT and HET genotypes combined) and *nee* mice. Representative images are shown of an optic nerve cross-section with myelin staining by paraphenylenediamine (PPD) from 8-week-old **(D)** WT and **(E)** *nee* mice. Note the decreased axon density and increased PPD axoplasm staining in panel E compared to panel D. Scale bar = 10 μm. Plots of **(F)** optic nerve cross-sectional area and **(G)** optic nerve total axon counts showing a statistically significant differences between 8-week-old WT and *nee* mice. **p < 0.01; ***p < 0.001

### 3.2 Nuclear size characteristics of surviving BRN3A^+^ RGCs decrease with increasing severity of glaucomatous damage in nee mice

Using the large amount of systematically collected feature data for the 748,782 segmented retinal nuclei underlying our study of *nee* mice, we examined whether any of the feature parameters automatedly measured by ImageJ correlated to disease stage. Region of interest feature data captured by ImageJ was averaged for each retina and tested for correlation to BRN3A^+^ cell density (Fig. 3). Three populations were considered: WT controls, *nee* retinas with no or mild disease (*nee*^*norm*^; within 2 SD of WT density of BRN3A^+^ nuclei), and *nee* retinas with moderate to severe disease (*nee*^*mod-sev*^; < 2 SD of WT density of BRN3A^+^ nuclei).

**Fig. 3.**
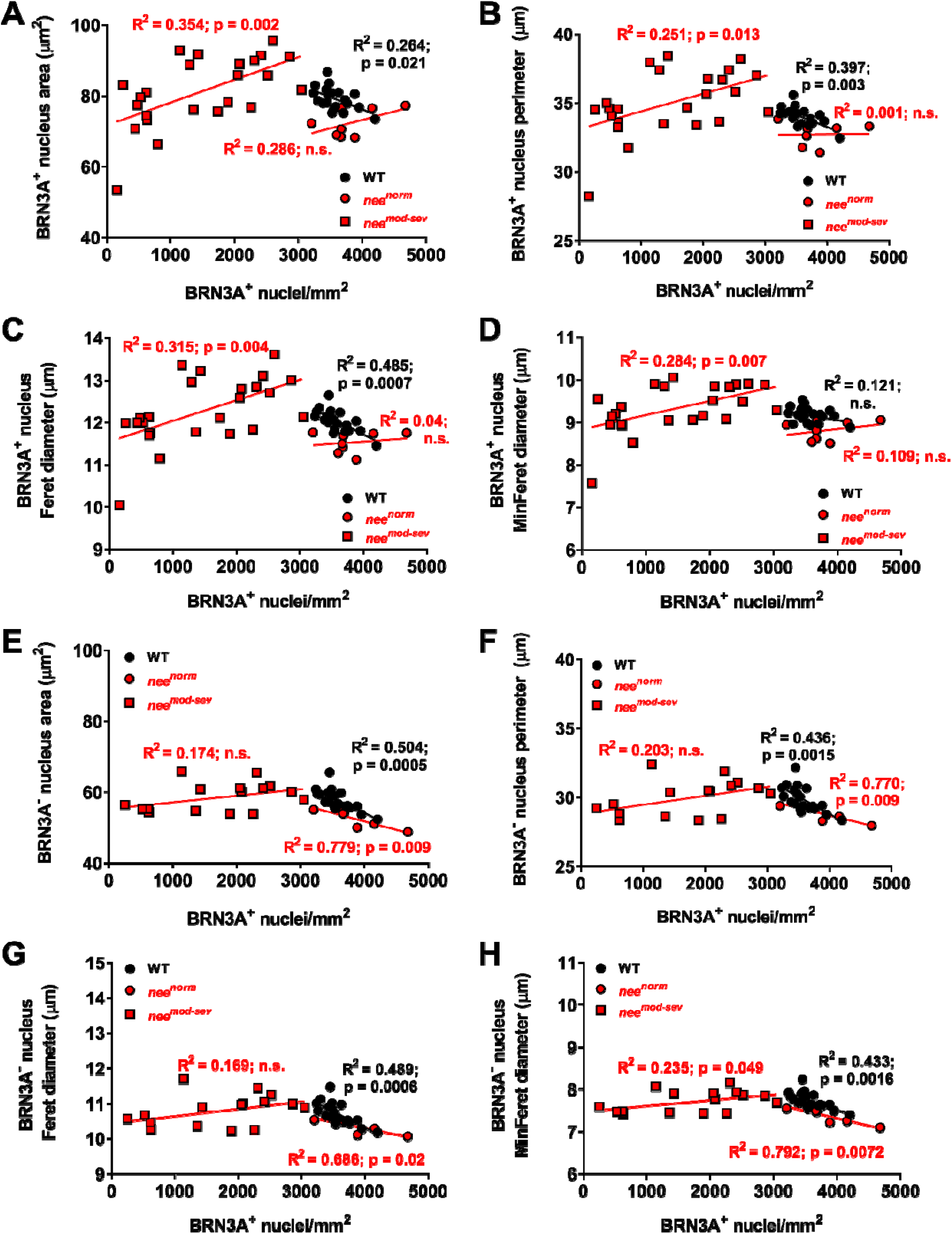
Correlations between automatedly collected nuclei density and size metrics for BRN3A^+^ and BRN3A^−^ nuclei. Average BRN3A^+^ RGC nuclear **(A)** area, **(B)** perimeter, **(C)** Feret diameter, and **(D)** MinFeret diameter plotted as a function of average density for individual *nee* and WT retinas. The BRN3A^+^ RGC nuclei of *nee*^*mod-sev*^ retinas consistently showed significant trends that nuclei size characteristics decrease with increasing severity of glaucomatous damage. Average BRN3A^−^ nuclear **(E)** area, **(F)** perimeter, **(G)** Feret diameter, and **(H)** MinFeret diameter plotted as a function of average density for individual *nee* and WT retinas. Unlike BRN3A^+^ RGC nuclei, the nuclei of *nee*^*mod-sev*^ retinas did not robustly show trends that nuclear size characteristics change during glaucoma. WT *(black circles)*; *nee*^*mod-sev*^ (*red squares*), diseased *nee* retinas with BRN3A^+^ nuclei density < two SD compared to WT; *nee*^*norm*^ (*red circles*), healthy *nee* retinas with BRN3A^+^ nuclei density comparable to WT.

Among BRN3A^+^ RGCs (Fig. 3A-D), WT retinas showed a significant correlation between average nuclear area and density (Fig. 3A) in which average nuclear size slightly increased as density decreased across the normal range of densities (3,238-4,203 nuclei/mm^2^; mean = 3,577, 1 SD = 241.5). Contrasting this, the RGCs of *nee*^*mod-sev*^ retinas showed an opposite trend that was also significant, in which nuclei became progressively smaller on average as density fell during disease. RGCs of *nee*^*norm*^ retinas had no detectable correlation between nuclear area and density. Similar trends and statistical associations were detected in plots of BRN3A^+^ density versus nuclear perimeter (Fig. 3B), Feret diameter (Fig. 3C), and MinFeret diameter (Fig. 3D). Among BRN3A^−^ nuclei (Fig. 3E-H), WT retinas again showed a significant correlation between average nuclear area and density (Fig. 3E) in which BRN3A^−^ nuclei became slightly larger with decreasing density of BRN3A^+^ nuclei. Unlike RGCs, the BRN3A^−^ population of nuclei in *nee*^*mod-sev*^ retinas showed no significant correlation to density. Also differing from RGCs, the BRN3A^−^ population of nuclei in *nee*^*norm*^ retinas mirrored the trends of WT cells, with average nucleus size slightly increasing as density decreased. Similar trends and statistical associations were detected in plots of BRN3A^−^ density versus nuclear perimeter (Fig. 3F), Feret diameter (Fig. 3G), and MinFeret diameter (Fig. 3H), with the sole exception of one association between BRN3A^−^ density and MinFeret diameter in *nee*^*mod-sev*^ retinas that met statistical significance with a slim margin (p = 0.049). In sum, these results point to two events influencing nuclear size. In almost all analyses (regardless of genotype or marker) there was a slight enlargement of nuclei as RGC density decreased. At approximately 3,000 nuclei/mm^2^ there was an inflection point in the trend lines, below which only the population of viable RGC nuclei, but not BRN3A^−^ cells, had nuclei that became smaller as disease was progressively severe.

### 3.3 RGCs with small nuclei account for a greater proportion of the surviving population in severe stages of nee glaucoma

We next sought to explore what phenomena might be causing the average nuclear size to be decreasing through disease progression. First, we analyzed the population distribution of BRN3A^+^ nuclear size in *nee*^*mod-sev*^ mice compared to WT littermates (Fig. 4). When all retinas were grouped together, the average size of BRN3A^+^ nuclei in *nee*^*mod-sev*^ retinas was 6.8% larger (84.5 ± 21.5μm^2^) than WT (79.1 ± 18.3μm^2^; p < 1E10^−15^; Fig. 4A) presumably because many retinas with less severe RGC loss had an average nuclear size that was larger than WT (Fig. 4A, B). However, a limitation of combining data in this way was that the healthiest *nee*^*mod-sev*^ retinas, those with the greatest number of cells, were overrepresented. Thus, instead of studying a size distribution for all retinas of a genotype together, we next analyzed distributions for each retina individually.

**Fig. 4.**
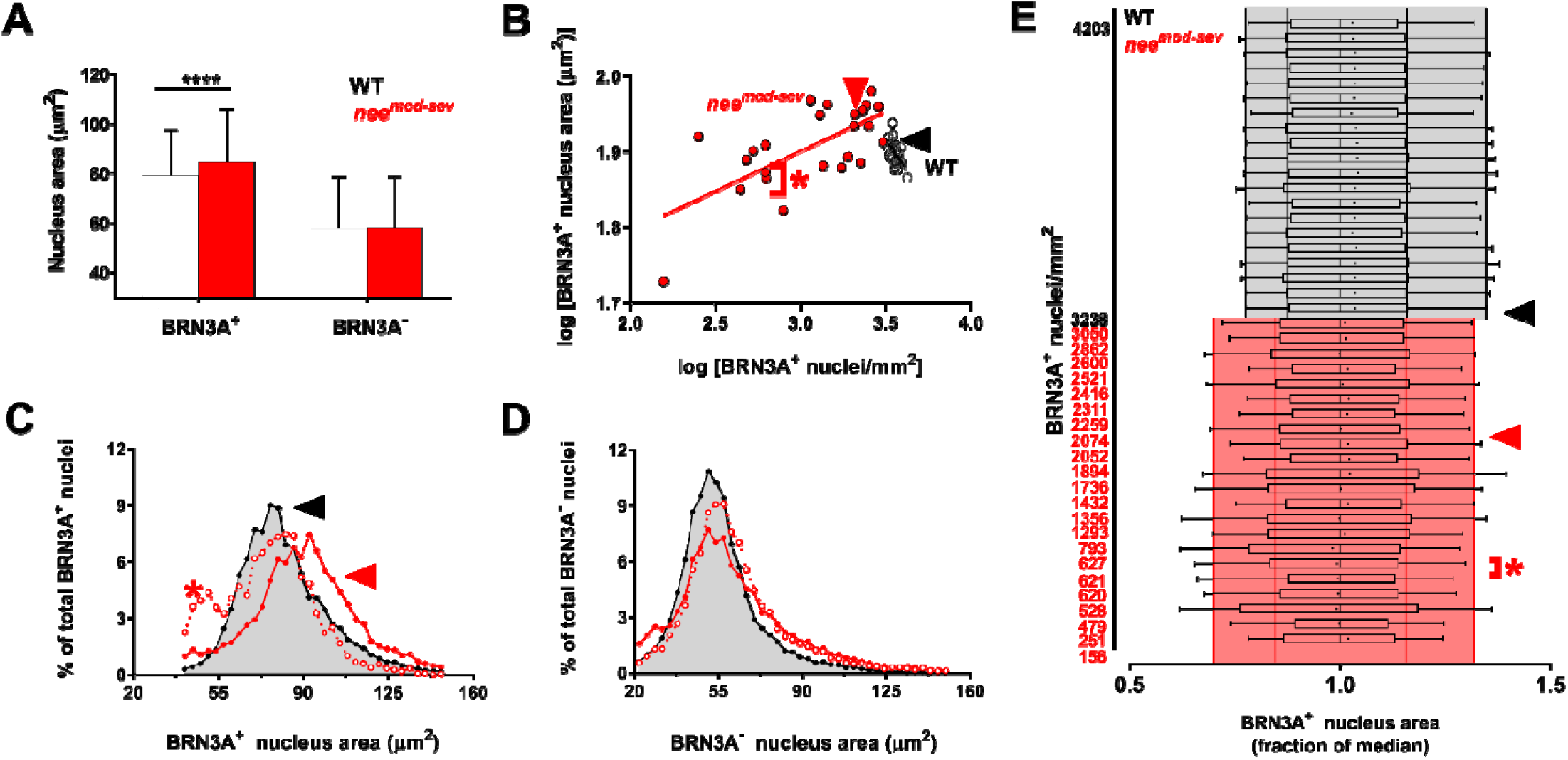
Population distributions of nuclei size in *nee*^mod-sev^ retinas. **(A)** Average area of nuclei with data from all retinas aggregated by genotype. **(B)** Average area of nuclei with each retina individually plotted. Surviving BRN3A^+^ RGC nuclei in *nee*^mod-sev^ retinas (red) were on average larger than WT (black). Note that many retinas with relatively modest RGC loss had an average nuclear size that was larger than WT; as glaucoma progressed the average size decreased. **(C)** Representative distribution curves for BRN3A^+^ nuclei, showing a WT retina (black arrowhead), a *nee* retina with modest disease (red arrowhead), and a *nee* retina with severe disease (red asterisk, in this instance data were combined from both retinas of a *nee* mouse in which both retinas had similar damage, increasing the number of nuclei analyzed to a number similar to the other examples). The same representative retinas are similarly indicated in panels B, C, and E. Note that in modest disease, the distribution curve maintained a similar shape compared to WT, but was shifted to the right, indicating nuclear enlargement broadly occurring in most RGCs regardless of their initial size. In severe disease, the right-hand tail of the curve was attenuated and there was a corresponding reciprocal peak in the left-hand portion of the curve, indicating an increased proportion of small RGC nuclei relative to the total population. **(D)** Distribution curves for BRN3A^−^ nuclei. **(E)** Box and whisker plot for each WT (grey) and *nee*^mod-sev^ (red) retina. To highlight differences in the shape of the distribution, data are normalized to the median value of nucleus size for each retina. Mean (black dot), median (solid line within each box), and percentile values (10th, 25th, 75th and 90th; vertical lines, black for WT and red for *nee*^mod-sev^) relative to the median. The average range of the distribution between the 10th and 90th percentiles (shaded background) was displaced leftward in *nee*^mod-sev^ relative to WT.

Retinas from WT mice consistently exhibited a broad distribution of BRN3A^+^ nuclear size with a preponderance of small nuclei but a long right-hand tail (Fig. 4C, E). In these retinas, the average BRN3A^+^ nuclear size was larger than the median in every case. An example of such a retina is indicated with a *black arrowhead* in Fig. 4 B&C, in this instance a retina with 3,238 BRN3A^+^ nuclei/mm^2^ (Fig. 4B) having a long right-hand tail (Fig. 4C), with average and median nuclear sizes of 80.9 ± 18.1 μm^2^ and 78.1 μm^2^, respectively.

In contrast, the size distributions were highly variable among *nee*^*mod-sev*^ retinas. Some *nee*^*mod-sev*^ eyes with comparatively modest RGC loss had a rightward shift in the entire histogram, whereby there was an increase in both the median and the mean relative to WT, with minimal change in the shape of the curve. An example is illustrated by the *red arrowhead* in Fig. 4 B&C. If the greater average nuclear size were caused by a preferential loss of smaller nuclei, the left part of the curve would decrease, and the right side would reciprocally increase (note the Y axis represents the percentage of total BRN3A^+^ nuclei). Because the shape of the histogram of this modestly affected eye only changed minimally while the entire curve was shifted rightward, the increased average size was not a result of preferential loss of the smallest cells and was more likely reflective of nuclear enlargement broadly occurring in most RGCs regardless of their initial size. In severe disease, the shape of the histogram was markedly different. Severely affected *nee*^*mod-sev*^ eyes had fewer large nuclei and a larger proportion of smaller RGCs. An example is illustrated by the *red asterisk* in Fig. 4 B&C (in this instance, combining data from two severely affected retinas, from a single mouse, with very similar BRN3A^+^ densities—so that a larger number of nuclei, similar to the other comparisons, could be considered). The right-hand tail of the curve was attenuated and there was a corresponding reciprocal peak in the left-hand tail of the curve, representing an increased proportion of small RGC nuclei relative to the total population. In this case, the smaller average nuclear size in severely affected eyes reflected a greater proportion of small nuclei in the surviving RGC population.

To test whether these differences in the nuclear size distribution were specific to RGCs, we plotted histograms of BRN3A^−^ nuclear size quantified from the same eyes and did not observe as pronounced differences in histogram shape or position of the major peak (Fig. 4D). As expected for a population likely consisting mostly of displaced amacrine cells (Drager and Olsen, 1981; Janssen et al., 2013), the average size of BRN3A^−^ nuclei was smaller than BRN3A^+^ nuclei.

To broadly visualize how the shape of the nuclear size distributions related to disease state, distributions were analyzed individually for all retinas and graphed in a box and whisker plot (Fig. 4E). In this representation, every retina was normalized to its median to allow for comparisons of the shape of the distribution. In WT retinas, the mean value was consistently greater than the median, reflecting the long right-hand tail of the distribution. In contrast, many *nee*^*mod-sev*^ retinas exhibited a smaller difference between the median and the mean, where the mean was displaced leftward towards the median, suggesting an attenuated right-hand tail of the curve and a greater proportion of smaller nuclei compared to the WT cell population. In some *nee*^*mod-sev*^ retinas, the mean was displaced even farther to the left beyond the median, signifying a longer left-tailed distribution with proportionately more RGC nuclei of small size. The greater proportions of small nuclei in *nee*^*mod-sev*^ retinas was also indicated by a decrease in the average of the 10^th^, 25^th^ and 90^th^ percentile values of *nee*^*mod-sev*^ retinas compared to WT.

## 4. Discussion

In this study with glaucomatous *nee* mice, ImageJ was used to quantify some traditional phenotypes commonly needed in studies of glaucoma (density of variously defined cell populations over time), as well as some novel phenotypes (size metrics for automatedly detected regions of interest) whose relevance to glaucoma was unknown. The methodology for RGC nuclei quantification has some elements of advancement within it, but the focus here has been on its application.

The first component of the study was to study the time course of neurodegeneration in *nee* retinal flat-mounts over time. These experiments indicated that *nee* mice have a 49% reduction in BRN3A^+^ RGCs by 8 weeks of age. No eyes escaped disease; every retina from mice aged 8 weeks or older was affected. However, some cases of severely affected retinas were already detectable at 4 weeks of age (our youngest cohort). While the early onset and complete penetrance of glaucoma are advantage of the *nee* model for testing neuroprotective therapies, it is important to recognize that the model does have some heterogeneity in age of onset and treatments would presumably need to be initiated early. Based on the current findings, initiation of treatments in newborn mice no later than 4 weeks of age appears warranted, and in some cases, treating pregnant/nursing mothers may be advantageous. From the BRN3A^+^ cell densities measured in 8-week-old WT and *nee* mice, we can predict that a neuroprotection of 50% of the RGCs otherwise expected to lose BRN3A marker expression and/or die from glaucoma should be detectable with 90% power (α = 0.05) using a two-tailed *t*-test with cohorts of 11 samples per treatment group. From the RGCC thickness measured in 8-week-old non-mut and *nee* mice, we can predict that RGCC thickness increases of 50% of the thickness lost in *nee* mice should be detectable with 90% power (α = 0.05) using a two-tailed *t*-test with cohorts of 18 mice per sample group. From the optic nerve cross-sectional area measured in 8-week-old WT and *nee* mice, we can predict that optic nerve cross-sectional area increases of 50% of the thickness lost in *nee* mice should be detectable with 90% power (α = 0.05) using a two-tailed *t*-test with cohorts of 18 mice per sample group. From the optic nerve axon counts quantified in 8-week-old WT and *nee* mice, we can predict that optic nerve axon count increases of 25% of the number of axons lost in *nee* mice should be detectable with 90% power (α = 0.05) using a two-tailed *t*-test with cohorts of 8 mice per sample group. Thus, tests of neuroprotective agents for *nee*-mediated early onset glaucoma are feasible for this model by quantifying BRN3A^+^ cell density, using OCT to measure RGCC thickness, measuring optic nerve cross-sectional area, and quantifying optic nerve axon numbers, with BRN3A^+^ cell density and optic nerve axon counts requiring fewer mice and directly quantifying RGC loss. Since a subset of RGCs in the *nee* model lose BRN3A expression prior to cell death, it is important to note that this power calculation is specific to use of BRN3A to mark RGCs. Use of other RGC markers may give different results depending on if or when RGCs lose marker expression prior to cell death. Quantification of cells that are nuclear-marker positive and RGC-marker negative, as well as use of H&E to count all retinal nuclei, can help parse loss of marker expression versus cell death as we have done here for the BRN3A RGC marker. Measurement of the RGCC via OCT requires more mice per cohort and is an indirect measurement of RGC loss but is a non-invasive assay that has the benefit of providing same-animal longitudinal data. Therefore, our current data should aid in the design of meaningful future experiments utilizing the *nee* model of early onset glaucoma.

An additional insight gained through these experiments was that RGC quantification data obtained from *nee* using immunolabeling techniques should be interpreted with caution. In our study, we detected an appreciable loss of immunoreactivity in RGCs. Approximately 20% of RGCs in *nee* mice aged 4–15 weeks lost BRN3A expression prior to cell death. This finding aligns with previous reports of reduced gene expression in models of acute RGC injury (Huang et al., 2006; Schlamp et al., 2001) and in DBA/2J mice (Soto et al., 2008). Accordingly, the 49% reduction in RGC density detected in the 8-week-old *nee* cohort was probably an overestimate of RGC loss—assuming 20% of the nuclei likely lost BRN3A immunoreactivity, the true degree of RGC death was likely closer to 30%. Importantly, no loss of BRN3A^−^ nuclei was observed in *nee* retinas. Sparing of non-RGC types is supportive of *nee* as a model of glaucoma, as opposed to other types of retinal neurodegeneration not specific to RGCs. Many studies with mouse models of glaucoma do not report the propensity for neuronal damage to non-RGC cell-types in the inner retina, such as displaced amacrine cells (Pang and Clark, 2020), but the specificity of RGC loss observed in *nee* mice do distinguish it from models based upon NMDA excitotoxicity (Hama et al., 2008; Siliprandi et al., 1992) or retinal ischemia (Dijk and Kamphuis, 2004; Ju and Kim, 2011) in which profound loss of other cell-types is known to occur.

Several mouse models relevant to early onset developmental forms of glaucoma have been described (Akula et al., 2020; Chang et al., 2001; Chen and Gage, 2016; Ishibashi et al., 1999; Iwao et al., 2009; Kroeber et al., 2010; Labelle-Dumais et al., 2020; Lachke et al., 2011; Smith et al., 2000; Thomson et al., 2017; Tolman et al., 2021). Among them, many studies have focused on phenotypes of the anterior segment and there have been relatively few in depth studies of RGC damage. One exception, a detailed study by Daniel et al. (Daniel et al., 2019), utilized *nee* mice to study the susceptibilities of 4 different RGC sub-types to glaucomatous damage. The overall levels of RGC damage in the study by Daniel et al. were assessed with NeuN immunolabeling (manual counting of 6–8 retinal flat-mounts per age; 8 sample regions per retina, each 0.09 mm^2^; 4 central and 4 peripheral) and detected a steady age-related decline in RGC density with statistically significant differences as early as 30 days of age (2,349 ± 126 cell/mm^2^ in wild-type, which reduced to 1,785 ± 161 cells/mm^2^ by day 30 in *nee* homozygotes). In comparison to the current findings, which had different goals and used different methodology (BRN3A immunolabeling; automated segmentation of 3–7 retinal flat-mounts per age; 12 sample regions per retina, each 0.18 mm^2^; 4 central, 4 mid-peripheral, and 4 peripheral), both studies detected an early onset glaucomatous loss of RGCs but the study by Daniel detected a statistically distinguishable loss earlier. Several factors likely contributed to these slightly different findings. First, the current study used BRN3A immunolabeling, which we found is lost in some RGCs prior to actual cell death, perhaps confounding an ability to detect differences. Second, there were notable differences in total RGC densities, even among wild-type controls between the two studies (∼2,400 cells/mm^2^, Daniel; ∼3,600 cells/mm^2^, current study), hinting that the study by Daniel had a greater emphasis on peripheral sites where RGC densities are lowest.

The second component of the current study involved a discovery-based analysis of the large amount of automatedly collected region of interest feature data captured by ImageJ. At the outset, it was unclear whether subtle differences in these “phenotypes” would add any information to understanding of glaucoma. At least hundreds, if not thousands, of previous studies have undoubtedly qualitatively observed RGC nuclei and a few have measured nuclear areas. In studying a much larger number of nuclei than practical manually, at a micron-level which might surpass the ability of human observers to notice trends, the broadest question of the current analysis was “would image-based phenotyping detect anything new?”. The results did detect significant trends, pointing to two events influencing nuclear size. In almost all analyses (regardless of genotype or marker) there was a slight enlargement of nuclei as RGC density decreased. At approximately 3,000 nuclei/mm^2^ there was an inflection point in the trend lines, below which only the population of viable RGC nuclei, but not BRN3A^−^ cells, had nuclei that became smaller as disease was progressively severe.

One event influencing nuclear size appears to be a passive “spreading out” of nuclei as space allows. Spreading appears to have a predominant influence on nuclear area in retinas with densities above 3,000 nuclei/mm^2^. For most size-related parameters of BRN3A^+^ and BRN3A^−^ nuclei, among both WT and *nee*^*norm*^ retinas, there was a strong negative correlation between nuclear size and cell density, whereby average nucleus size increased with decreasing cell density. We speculate that under normal physiological conditions, neurons generally tend to be larger in retinas with lower cell densities because space is less confined. The only exceptions to this finding were a single lack of significance for BRN3A^+^ nuclei in WT retinas for MinFeret diameter, and all four of the comparisons for BRN3A^+^ nuclei in *nee*^*norm*^ retinas. Among the *nee*^*norm*^ retinas, the lack of association might be explained by simultaneously competing events, a passive spreading out as RGCs begin to be lost that tends to increase average nuclear size, offset by a preferential loss of larger nuclei that tends to decrease average nuclear size. The phenomenon of spreading appears to also be present in an anatomical context, where RGC size is known to vary with respect to eccentricity. RGCs are significantly larger in the peripheral retina, an area of relatively low cell density compared to the central retina, where RGCs are smaller and more numerous (Drager and Olsen, 1981; Urcola et al., 2006). RGC enlargement in some stages of disease has been previously observed in multiple studies (Davis et al., 2020; Hedberg-Buenz et al., 2016a; Moore and Thanos, 1996; Urcola et al., 2006). For example, among surviving RGCs in a rat model of chronic IOP elevation induced by microbead injection (Urcola et al., 2006), in which a loss of 27.2% of RGCs after 24 weeks led to a 11.9% increase in RGC somal area—a subtle change in size in response to modest changes in RGC density similar to those observed here. Urcola et al. described this as “hypertrophy”. Because RGC somal and nuclear areas have previously been shown to be positively correlated (Davis et al., 2020; Janssen et al., 2013), it seems likely that the “hypertrophy” observed by Urcola et al, and “spreading” described here, are the same phenomenon. Davis et al. (Davis et al., 2020) observed RGC nuclear enlargement in the early stages following partial optic nerve transection and hypothesized that it might involve metabolic responses of the injured RGCs. Because we have observed spreading in not only glaucomatous RGCs, but also in non-RGCs of mice with glaucoma and in RGCs of healthy WT mice, we propose a more passive response—though future studies to study this phenomenon are needed.

A second event influencing nuclear size appears to be a preferential loss of larger RGC nuclei through progressive stages of glaucoma. For only BRN3A^+^ nuclei of *nee*^*mod-sev*^ retinas, there was a strong positive correlation with RGC nuclear size, whereby average nucleus size decreased with decreasing cell density. This phenomenon was specific to RGCs and not detected among BRN3A^−^ nuclei. In examining size distribution plots, the RGC population in glaucomatous retinas was markedly different from WT retinas and highly dependent on disease stage. Healthy WT retinas consistently exhibited a right-skewed distribution of BRN3A^+^ nuclear size, in which the average RGC size was always greater than the median, and the right-hand side of the distribution curve had a long tail. In retinas with modest glaucoma, there was rightward displacement of the entire distribution curve, reflecting an indiscriminate enlargement of RGC nuclei of all size. In retinas with advanced glaucoma there was an attenuation of the right-hand tail of the size distribution, in some cases becoming a left-skewed distribution, and there was often a second peak in the left-hand tail, indicating an enlarged fraction of small RGC nuclei.

We speculate that the biological basis for greater proportions of small RGC nuclei in eyes with severe disease may involve nuclear atrophy and/or size-selective vulnerability to glaucoma. Some aspects of apoptosis are rapid events, with multiple studies identifying active apoptosis in only <1–15% of RGCs at any given time (reviewed in (Cordeiro et al., 2011); thus, many apoptotic events are likely too transient to significantly impact the population average for nuclear size. In contrast, nuclear atrophy driven by histone deacetylase activity is a sustained event that can precede BAX-dependent apoptosis in RGCs (Janssen et al., 2013; Schmitt et al., 2016). In mouse models of optic nerve crush, nuclear atrophy appears within 24 hours, is maximal at 5 days, and persistent thereafter (Janssen et al., 2013). In comparison to other events occurring with optic nerve crush, nuclear atrophy precedes both apoptotic nuclear fragmentation, which typically begins at days 5–7 post-injury, and cell death, which typically begins after day 10 post-injury (Li et al., 1999). Thus, nuclear atrophy is a sustained event that could contribute to our current observations with *nee* mice. Alternatively, average RGC size could decrease with progressive disease because the largest cells are preferentially dying. Preferential vulnerability of larger RGCs has been a long-standing hypothesis in glaucoma (Glovinsky et al., 1991; Vickers et al., 1995; Wang et al., 2020), and in mice, has been supported by an enrichment of small RGCs in old DBA/2J mice with late-onset glaucoma (Buckingham et al., 2008; Hedberg-Buenz et al., 2016a). The current results establish that this is also the case for *nee* mice with early onset congenital glaucoma.

There are caveats to consider in the interpretation of these data. First, immunolabeling was used to quantify only BRN3A^+^ RGCs because it labels the nucleus, lending itself nicely to the semi-automated high-throughput segmentation approach used here. Accordingly, the data most specifically relate to the surviving RGC population that still expresses BRN3A. Although *Brn3a* appears to be transcribed in all sub-classes of RGCs (Rheaume et al., 2018), most studies find that immunolabeling fails to identify some (∼3–15%) RGCs (Davis et al., 2020; Galindo-Romero et al., 2011; Rodriguez et al., 2014). Thus, RGCs that either lost, or never had, detectable BRN3A were potentially confounding. For example, many melanopsin containing RGCs are not well-labeled by BRN3A (Sanchez-Migallon et al., 2018; Valiente-Soriano et al., 2014). Because melanopsin-containing RGCs are relatively rare (∼2–3% of the total RGC population) (Valiente-Soriano et al., 2014), these cells likely had a miniscule impact on our overall findings. Non-RGC cell-types were only studied in aggregate, not with additional specific markers. This may be of relevance because a subtle loss of displaced amacrine cells that are coupled to RGCs via gap junctions has been reported in a microbead-induced model of glaucoma (Akopian et al., 2019). Our current experimental design, lacking specific markers of displaced amacrine cells and complicated by RGCs losing BRN3A expression, was not well-matched for detection of this event—though it would be an interesting area for future work using *nee* mice. Second, with 40+ sub-types of RGCs now molecularly recognized (Rheaume et al., 2018; Tran et al., 2019), it is probable that nuclear size may be a susceptibility-factor for many RGCs, but other sub-type specific factors could also play a role (Tran et al., 2019; Wang et al., 2020). Melanopsin-containing RGCs, which are somewhat resistant to multiple forms of damage (Cui et al., 2015; Daniel et al., 2019; Jakobs et al., 2005; Li et al., 2006; Sanchez-Migallon et al., 2018), are often described as an exception to the “larger = more susceptible” hypothesis, perhaps incorrectly, as their soma are actually small to average in size and only their dendritic field is comparatively large (Berson et al., 2010; Coombs et al., 2006; Xiao et al., 2021). In future work, it could be informative to elucidate the molecular events which drive the association of smaller nuclei with RGC survival.

In sum, this study has quantified the severe loss of RGCs in the *nee* mouse model of congenital glaucoma and used a large amount of simultaneously collected image data related to nuclear size to phenotype glaucomatous mice in a new way. The results point to competing events influencing nuclear size, with spreading of multiple cell types occurring with modest decreases in RGC density and a preferential loss of large RGC nuclei in moderate to severe cases of RGC loss due to glaucoma. Given the progress in refining approaches using automated image analysis to quantify RGC soma (Danias et al., 2006; Dordea et al., 2016; Guymer et al., 2020; Hedberg-Buenz et al., 2016a; Hedberg-Buenz et al., 2016b; Masin et al., 2021; Salinas-Navarro et al., 2009) and axons (Bosco et al., 2015; Mysona et al., 2020; Reynaud et al., 2012; Ritch et al., 2020; Teixeira et al., 2014; Zarei et al., 2016) it appears that the approach that we, and others (Davis et al., 2020), have begun to utilize to phenotype RGCs in these automated ways may make additional discoveries. Although studies to date have analyzed only a small number of parameters that were all related to nuclear size, future experiments could likewise analyze many additional features, some of which may also be beyond the practical limit for humans to have been led to suspect as important based only on visual observation.

## ACKNOWLEDGEMENTS

This work was supported in part by Merit Review Award (I01 RX001481) from the U.S. Department of Veterans Affairs RR&D Service and an NIH/NEI Center Support Grant to the University of Iowa (P30 EY025580). AHB was supported by Training Grant Award (T32DK112751). The funding sources had no involvement in study design; in the collection, analysis and interpretation of data; in the writing of the report; or in the decision to submit the article for publication. The contents do not represent the views of the U.S. Department of Veterans Affairs or the U.S. Government.

---

**Supplementary Fig 1.**
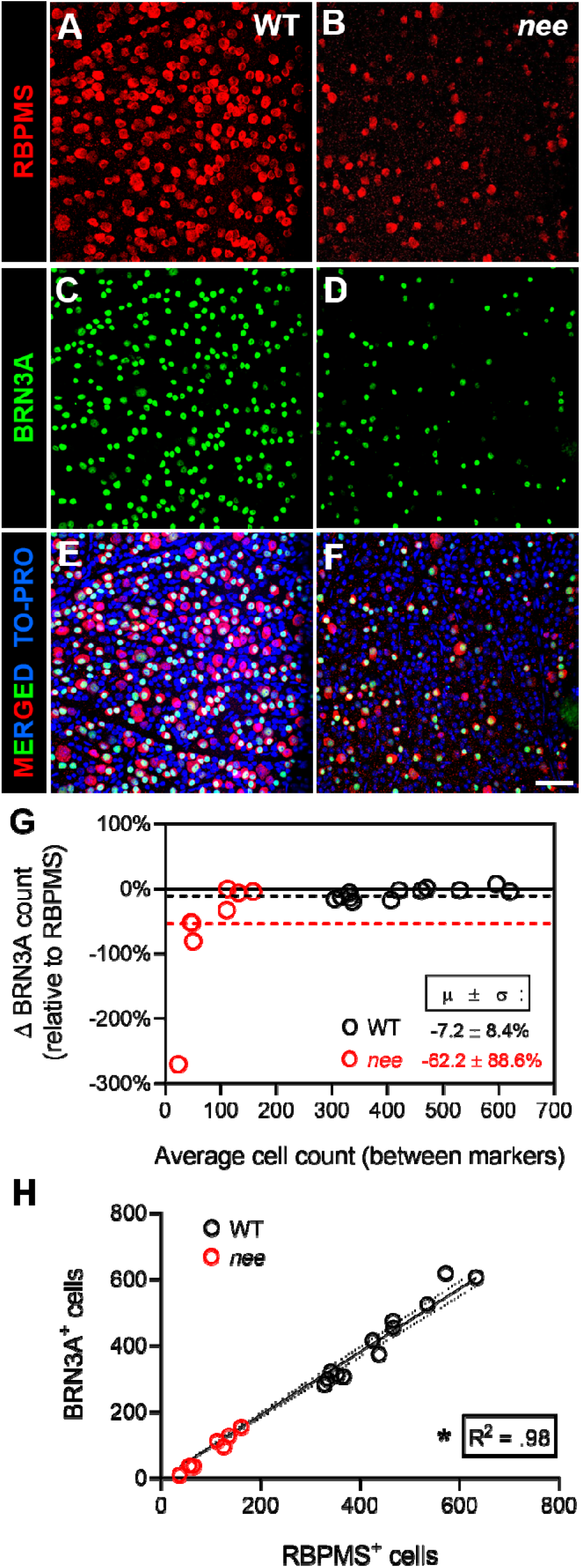
Co-labeling of retinas with BRN3A and RBPMS. Manual absolute cell counts of sampling field images captured from the peripheral retina immuno-labeled with two different markers for retinal ganglion cells (RGCs) to compare the proportion of mono- and dual-labeled cells in 6-wk-old *nee* vs. WT mice. Representative examples cropped from sampling fields of peripheral retina from a wild-type (WT, *left column*) and *nee* (*right column*) mouse with immunolabeling for **(A-B)** RBPMS (*red*) and **(C-D)** BRN3A (*green*), along with **(E-F)** nuclear staining by TO-PRO (*blue*) in composite view with merging of all channels. Scale bar = 50 μm. **(G)** Bland-Altman plot showing the percent (%) change in BRN3A^+^ absolute cell count of peripheral retina sampling fields (relative to RBPMS^+^) vs. the average absolute cell count of the two markers in the same sampling fields. Each dot represents a manual absolute cell count of a peripheral retina sampling field using both markers. For each genotype, the average % change in BRN3A^+^ absolute cell counts of sampling fields are indicated by inset dotted lines and equations (*average with standard deviation, μ ±* σ) with distinct coloring that corresponds to each genotype. **(H)** Graph plotting manual absolute cell counts of BRN3A^+^ vs. RBPMS^+^ cells of the same fields (*n* = 4 fields per retina) from retinas of *nee* (*n* = 2 mice) and WT controls (*n* = 3 mice). Each dot represents the absolute cell count using both markers for a single sampling field. Within the graph, the solid line represents the best fit line and the flanking dotted curves indicate the 95% confidence interval for the data points. The coefficient of determination (R^2^) gives the proportion of variation in absolute cell count in a given sampling field between the two markers. Overall, a total of *n* = 6,012 RBPMS^+^ (*nee*: *n* = 751 cells, WT: *n* = 5,261 cells) and 5,604 BRN3A^+^ cells (*nee*: *n* = 611 cells, WT: *n* = 4,993 cells) were counted across a set of *n* = 18 sampling fields from the peripheral retina (*nee*: *n* = 8 fields, WT: *n* = 12 fields).

